# *Cryptosporidium* modifies intestinal microvilli through an exported virulence factor

**DOI:** 10.1101/2025.02.25.639898

**Authors:** Elena Rodrigues, Mitchell A. Pallett, Lorian C. Straker, Tapoka T. Mkandawire, Katarzyna Sala, Lucy Collinson, Adam Sateriale

## Abstract

*Cryptosporidium* is a common intestinal infection of vertebrates and a significant threat to public health. Within the epithelial layer of the intestine, the parasite invades and replicates. Infected cells are readily detected under the microscope by the presence of elongated microvilli, particularly around the vacuole where the parasite resides. Here, we identify a family of *Cryptosporidium* virulence factors that are exported into the host cell during infection and localise to the microvilli. We examine the trafficking and function of the most highly expressed family member, MVP1, which appears to control the elongation of microvilli through engagement of host EBP50 and CDC42. Remarkably, this mechanism closely mirrors that of an enteropathogenic *Escherichia coli* virulence factor, MAP, which is also known to drive host microvilli elongation during infection. This highlights a unique instance where eukaryotic and prokaryotic virulence factors have convergently evolved to modulate host actin structures through a similar mechanism.

## Introduction

The intestinal parasite *Cryptosporidium* follows a faecal-oral route of transmission, and infections are often waterborne. This is largely due to the robust nature of the parasite’s oocyst form; a metabolically dormant life cycle stage that is resistant to normal water treatment methods such as chlorination. Globally, an estimated 9% of mortality in children under the age of 5 is attributed to diarrheal diseases^1^. *Cryptosporidium* is among the leading causes of severe diarrhoea in children, driving mortality but also morbidity in the form of malnutrition, which can lead to more permanent effects such as impairment of cognitive development and growth stunting^2,3^. To make matter worse, malnutrition also appears to be a risk factor for severe *Cryptosporidium* infections in children^4^. Even brief periods of malnutrition are suggested to be sufficient to significantly exacerbate parasite burden through suppression of host cell apoptosis and cell-turnover^4,5^.

When a *Cryptosporidium* oocyst is ingested, it responds to environmental cues, such as bile salts and temperature, to release motile forms of the parasite. These motile forms, known as sporozoites, then invade and replicate within epithelial cells that line the intestine of the host organism. While *Cryptosporidium* is an intracellular parasite, it remains outside the host cytoplasm, encased in a vacuole. The parasite creates this unique niche by enveloping itself within the host cell membrane at the apical surface of the epithelium, creating many ‘parasitophorous vacuoles’ that line the lumen of the intestine^6^. After several rounds of asexual replication, known as merogony, the parasite differentiates into a dimorphic sexual stage, producing male and female gametes^7^. Fertilization produces newly infectious oocysts that are shed into the environment to infect more hosts.

To ensure survival, intracellular parasites must be able to modify and manipulate their host cell. Within the Apicomplexa family of parasites, to which *Cryptosporidium* belongs, *Toxoplasma* (causative agent of toxoplasmosis) and *Plasmodium* (causative agent of malaria) are known to export hundreds of effector proteins into their host cells to facilitate this^8,9^. Exported effector proteins are known to participate in a variety of functions, including evasion of host immune responses, nutrient scavenging and structural modulation of the host cell^10,11^. Comparatively, very little is known regarding host-exported factors of *Cryptosporidium*. With *Cryptosporidium’s* relatively small genome of around 9Mb, it is thought to have very streamlined metabolic pathways with a greater dependence on host cell nutrients and metabolites, thus more exported proteins may be required for this parasite to survive^12,13^.

Extensive remodelling of the host cell actin cytoskeleton is something that has been previously described during *Cryptosporidium* infection, despite the mechanisms being largely unknown. This is seen in the production of actin rich structures between parasite and host cell during infection, often referred to as the ‘actin pedestal’, which is thought to strengthen parasite adherence^14^. Another prominent example of actin manipulation by the parasite is the length of host cell microvilli during infection^15–18^. Epithelial cells that harbour the parasite are known to have drastically elongated microvilli compared to that of non-infected bystander cells, a phenotype that appears to be conserved across vertebrates with intestinal *Cryptosporidium* infections. This phenotype of elongated microvilli is mirrored during infection with pathogenic *Escherichia coli*, which are known to drive host membrane protrusion around their site of attachment to intestinal epithelial cells^19,20^. The effector protein MAP, translocated into epithelial cells by the bacteria enteropathogenic *E. coli* (EPEC), has been reported to induce abnormal and elongated filopodia formation during infection. MAP is thought to be stabilized at the apical host cell membrane through interaction of its C-terminal PDZ-binding domain with the host scaffold protein EBP50. Once sequestered at the membrane MAP, which is a guanine nucleotide exchange factor (GEF) can activate the small GTPase CDC42 leading to the formation of membrane protrusions^20,21^.

Here we have used a combination of bioinformatics, genetic manipulation, and super-resolution microscopy to identify a family of host-exported and microvilli-associated *Cryptosporidium* proteins. We further characterise the most highly transcribed protein of the family, MVP1, a small, unstructured protein seen to be present within a subset of *Cryptosporidium* secretory organelles prior to invasion of the host epithelial cell. We establish that MVP1 is exported into the infected host cell during early stages of infection and remains present throughout the parasite lifecycle. Once inside the host, MVP1 localizes to the microvilli where it interacts with the host scaffold protein EBP50 via a conserved PDZ domain. Here MVP1 facilitates the elongation of the host microvilli, a phenotype that is lost during infection with an MVP1 deficient parasite line. Lastly, we highlight similarities between *Cryptosporidium* MVP1 and that of the bacterial effector MAP, which appear to have convergently evolved to manipulate the same host signalling pathways and cellular structure.

## Results

### cgd6_40 is a dense granule protein that is exported into the host cell early in *C. parvum* **infection**

To identify potential host-exported proteins of *Cryptosporidium,* we used an in-silico approach that made use of publicly available genomic and transcriptomic data sets^22^. We selected genes based on three criteria: firstly, on the presence of a predicted N-terminal signal peptide which suggests the protein to be secretory. Secondly, to look for proteins with high genetic variation and thus more likely to be exposed to the host cell immune system, we screened for genes with a high non-synonymous to synonymous ratio (dN/dS > 2). Lastly, genes were chosen based on high RNA expression early in infection, during asexual and sporozoite stages (the top 99^th^ percentile). When all three of these criteria were satisfied only one suitable candidate was identified, this was the gene cgd6_40 at the subtelomeric region of chromosome 6 (Figure 1a). Using CRISPR/Cas9-driven homologous recombination we fused a three-hemagglutinin epitope tag (3xHA) to the C-terminus of endogenous cgd6_40 within *C. parvum*. Downstream of this, and under a sperate promoter, we inserted a NanoLuciferase reporter for transgenic detection fused to a neomycin resistance marker for transgenic selection (Figure 1a). Transgenic parasites were selected for *in vivo* using *Ifn*γ*^-/-^* mice, as the parasite is not able to be effectively propagated within cell culture. NanoLuciferase output was measured from faecal samples taken daily as a proxy for parasite burden (Figure 1a). With such high transcription of cgd6_40 early in the parasite life cycle, we first looked for protein expression within sporozoite stages. Immunofluorescence staining at these stages with our tagged strain showed a strong HA expression with newly excysted sporozoites showing several loci towards the centre of the parasites. To resolve the localization in more detail we employed expansion microscopy coupled with super resolution microscopy^23^. HA staining in the sporozoites coincided with a recently described subset of secretory organelles within the parasite, known as small granules ^24^ (Figure 1b). In addition to this, it was also surprising to see strong HA staining present in a ring around the tip of a second organelle – the rhoptry.

**Figure 1.**
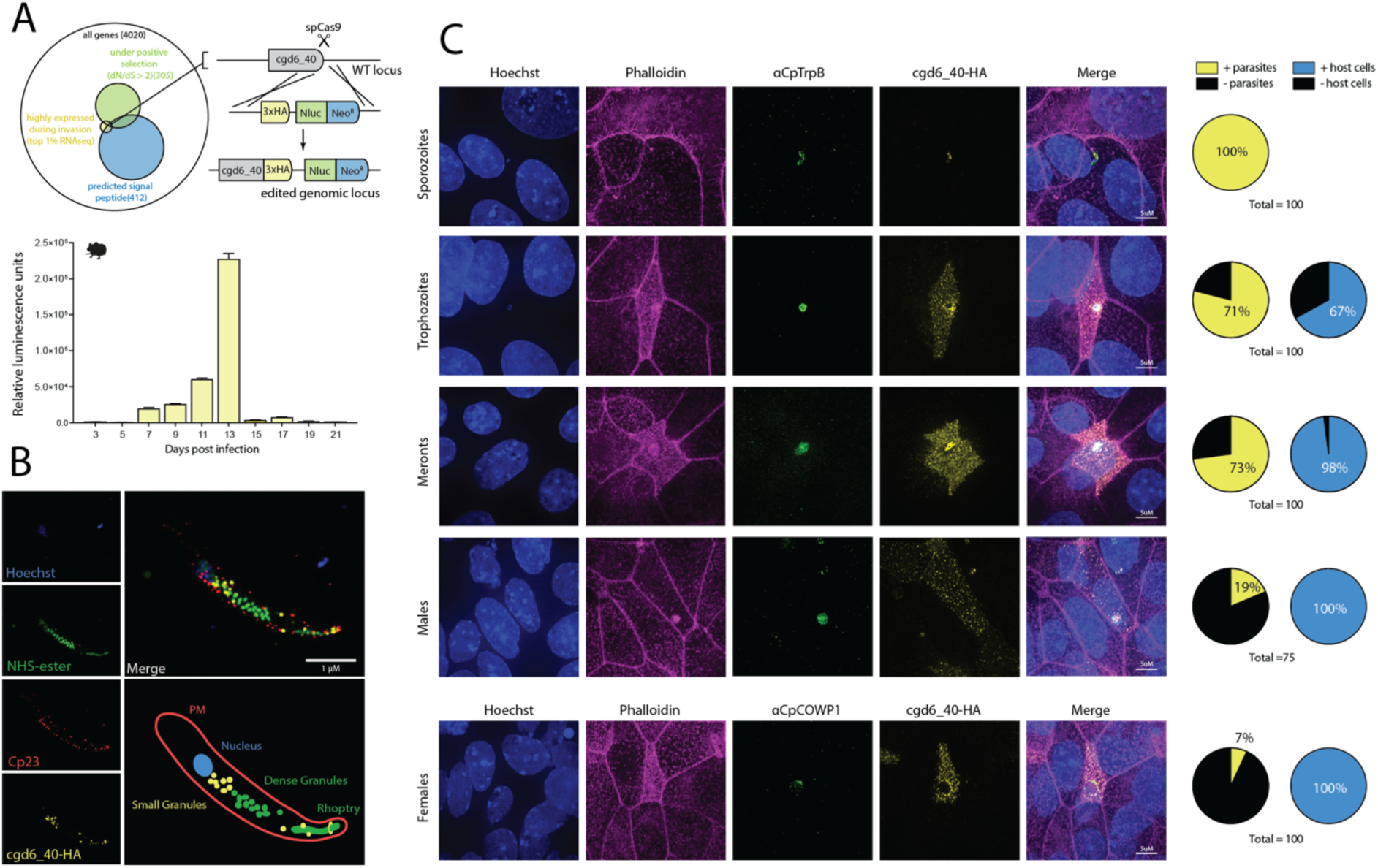
Cryptosporidium parvum cgd6_40 is trafficked into the infected host cell during infection. **A)** Diagram of the selection process of cgd6_40. Criteria included: (1) a predicted signal peptide (2) a high non-synonymous/synonymous ratio (2>dN/dS) and (3) highly expressed at sporozoite and asexual life cycle stages within publicly available transcriptome data sets (top 1% of RNAseq by tpm). Schematic for epitope tagging cgd6_40 at the genetic locus along with insertion of constitutively expressed NanoLuciferase reporter and neomycin resistance marker. Graph showing the relative luminescence reading from collected fecal samples during the initial transfection and selection of cgd6_40-HA transgenic parasites within *Ifn*γ*^-/-^* mice. Mean and standard deviation for three technical repeats. **B)** Expansion microscopy of a sporozoite with epitope labelled cgd6_40. Within sporozoites, HA-tagged cgd6_40 can be readily identified within a subset of dense granules known as the small granules^24^ **C)** IFA epithelial cell (HCT-8) monolayers infected with transgenic cgd6_40-HA parasites. Infections were fixed at 2.5 minutes (sporozoites), 6 hours (trophozoites), 24 hours (meronts) and 48 hours (males and females). Quantification of MVP1-HA protein localisation during infection at different life cycle stages on right.

Next, we infected ileocecal colorectal adenocarcinoma (HCT-8) cells grown in a monolayer with cgd6_40-HA parasites, using immunofluorescence to localize the tagged protein (Figure 1c). Cgd6_40-HA was present within all sporozoites prior to host cell contact and could be detected by immunofluorescence within host cells at roughly 3 hours into infection (Supplementary Figure 1a). As parasites matured from trophozoites to meronts, cgd6_40-HA appeared to accumulate within the infected host cell and when newly formed merozoites emerged to invade new cells, these parasites also contained loci of cgd6_40-HA (Supplementary Figure 1b). This suggests that the motile forms of the parasites, sporozoites and merozoites, both contain cgd6_40 prior to invasion and then continue to synthesize and export this protein during replication. Sexual stages, both male and female, were largely negative for cgd6_40-HA loci within the parasite yet tagged protein could be readily detected still present within their respective infected host cells (Figure 1c).

### cgd6_40 localizes intracellularly to the host microvilli apical membrane during *C. parvum* **infection**

Having determined cgd6_40 localization during parasite growth and differentiation, we next sought to confirm that cgd6_40 is exported into the cytoplasm of the infected epithelial cell. Monolayers of intestinal epithelial (HCT-8) cells were infected with cgd6_40-HA parasites and either permeabilized or not, following fixation. HA tagged protein was not detectable by immunofluorescence within infected cells that were not permeabilized, indicating that cgd6_40 is an intracellular protein (Figure 2a). Cgd6_40 was originally selected based on the presence of a predicted signal peptide which suggests that the protein enters a secretory pathway within *Cryptosporidium* prior to being exported into the host cell. To test this, we used Brefeldin A, which is known to inhibit protein transport from the endoplasmic reticulum to the Golgi, blocking the secretory pathway. Brefeldin A treatment of epithelial cells during infection with cgd6_40-HA parasites ablated cgd6_40-HA export into the host cells and resulted in an accumulation of protein within the parasites, supporting the prediction that cgd6_40 enters the host cell through the *Cryptosporidium* secretory pathway (Figure 2b). Subsequently, we used super-resolution microscopy to focus on determining a more exact localization of the protein following export. Cgd6_40-HA transgenic parasites were used to infect epithelial cell monolayers, which were fixed for IFA at 24 hours into infection. Super resolution imaging along the z-axis of infected cells revealed that cgd6_40-HA colocalizes to actin filaments within the host cell microvilli on the brush border (apical side) of the intestinal epithelial cell (Figure 2c). We next wanted to establish whether cgd6_40 export and localization occur in a similar manner within a more physiologically relevant setting. Within the ileum of *Ifn*γ*^-/-^*mice, cgd6_40-HA is host-exported and is also visible within the brush border of the intestine (epithelial cell microvilli) (Figure 2c). Due to this specific localization of cgd6_40, we will refer to this protein, from here on, as Microvilli Protein 1 (MVP1).

**Figure 2.**
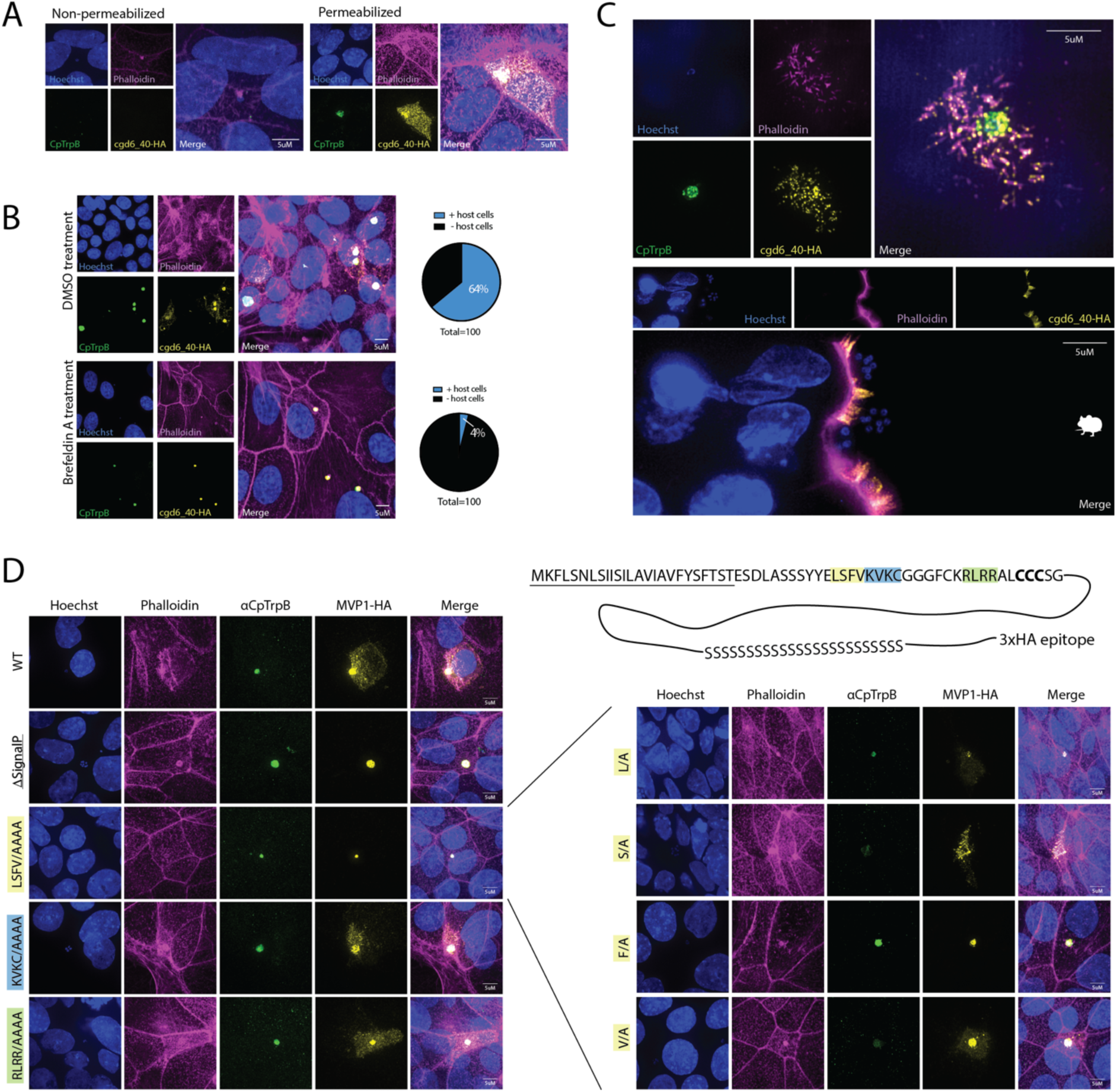
Cgd_40 relies on ER to Golgi transport and localises to the microvilli inside infected epithelial cells. **A)** Infection of epithelial cell (HCT-8) monolayers by cgd6_40-HA transgenic *C. parvum* parasites. Infections were fixed and either not permeabilized (left) or permeabilized (right) prior to staining. **B)** Epithelial cell monolayers infected with transgenic MVP1-HA *C. parvum* parasites and treated with Brefeldin A, which blocks ER to Golgi transport. Quantification of MVP1-HA protein localisation included on right. **C)** Super resolution microscopy MVP1-HA localisation *in vitro* (HCT-8 cell monolayer at 24 hours post infection, TOP) and *in vivo* (small intestine from *Ifn*γ*^-/-^* mice at day 10 post infection, BOTTOM). **D)** Epithelial cell monolayers infected with wild-type parasites transfected with plasmids driving altered MVP1-HA. Cell monolayers were fixed and labelled at 24hrs post infection. Note that removal of the predicted signal peptide and mutation of the conserved region ‘LSFV’ both caused a block in host export. Further point mutations demonstrated that mutation of phenylalanine alone, F39A, was sufficient to disrupt trafficking into the infected host cell.

### MVP1 relies on an essential phenylalanine to traffic into the host cell

Apicomplexa such as *Plasmodium* and *Toxoplasma* have well defined pathways for the trafficking and export of effector proteins. The canonical pathways for both organisms involve the recognition and cleavage of a specific amino acid motif by aspartyl proteases present within the secretory pathways, marking the protein for downstream processing and export. These cleavage motifs are termed the *Plasmodium* export element (PEXEL) motifs in *Plasmodium* and *Toxoplasma* export element (TEXEL) motifs in *Toxoplasma*^25,26^. As the host-export pathway for *Cryptosporidium* is still undescribed, we were interested to test whether MVP1 may also have an ‘export motif’ required for trafficking into the host cell. Within sequence alignments of the MVP1 gene from different intestinal *Cryptosporidium* species and strains, we observed that the genetic variation was largely present within the C-terminus of the protein, while the N-terminus showed a higher level of conservation between strains and species. Within the N-terminus we identified several highly conserved regions just downstream of the signal peptide and sought to determine whether any of these motifs played a role in trafficking into the host cell (Figure 2d). To accomplish this, we transiently expressed MVP1 mutants during infection, where either the signal peptide was removed or segments of conserved amino acids were changed to alanine residues. As expected, truncation of the signal peptide caused MVP1-HA to become trapped within the cytosol of the parasite, while full length MVP1 was freely exported into the host cell and accumulated within the microvilli. Mutation of the KVKC and RLRR conserved loci had no effect on MVP1 export or localization, however, mutation of the LSFV motif blocked the export from parasite to host cell. To dissect this further, an additional 4 mutants were created with single mutations of either the leucine, serine, phenylalanine, or valine residues within this LSFV motif (Figure 2d). Of all mutations, only phenylalanine mutated to an alanine residue (F39/A) blocked export of MVP into the host cell. This result was contrasting to the known *Cryptosporidium* host-exported protein MEDLE2 where a leucine residue (L35) was previously determined to be essential^27^. However, it was noted that L35 of MEDLE2 and F39 of MVP1 both fall to the right of a conserved serine, creating an analogous four amino acid motif that consists of a hydrophobic residue, a serine, and two further hydrophobic residues.

### MVP1 is part of a family of exported proteins that localise to the host microvilli

Although there were no easily identifiable homologs of MVP1, we noticed that within the *Cryptosporidium parvum* genome there were several small, unstructured proteins shared the same basic composition, with a predicted signal peptide, a similar export motif to MVP1, and a region of C-terminal serine repeats (Figure 3a and Supplementary Figure 2). Most of these proteins localized to a region on chromosome 2, previously thought to contain mucin-like glycoproteins. (Figure 3a). To look at these proteins in more detail, expression plasmids were created with each gene under the MVP1 promoter. These plasmids were transfected into *Cryptosporidium* parasites and localised at 24 hours into infection. Cgd2_390, cgd2_400, cgd2_410, cgd2_450 and cgd4_10 all appear to export and localize to the host microvilli and for this reason we chose to include them in the MVP family of proteins (Figure 3b and Supplementary Figure 2). Interestingly, all proteins that could be readily detected within the host cell contain a predicted palmitoylation site just downstream of the signal peptide. Genes lacking a predicted palmitoylation site (cgd2_420, cgd2_430 and cgd2_440), though sharing the other features common to the MVP family, could not be detected in the host cell by microscopy and could only be seen within the parasitophorous vacuole during infection (Figure 3b).

**Figure 3.**
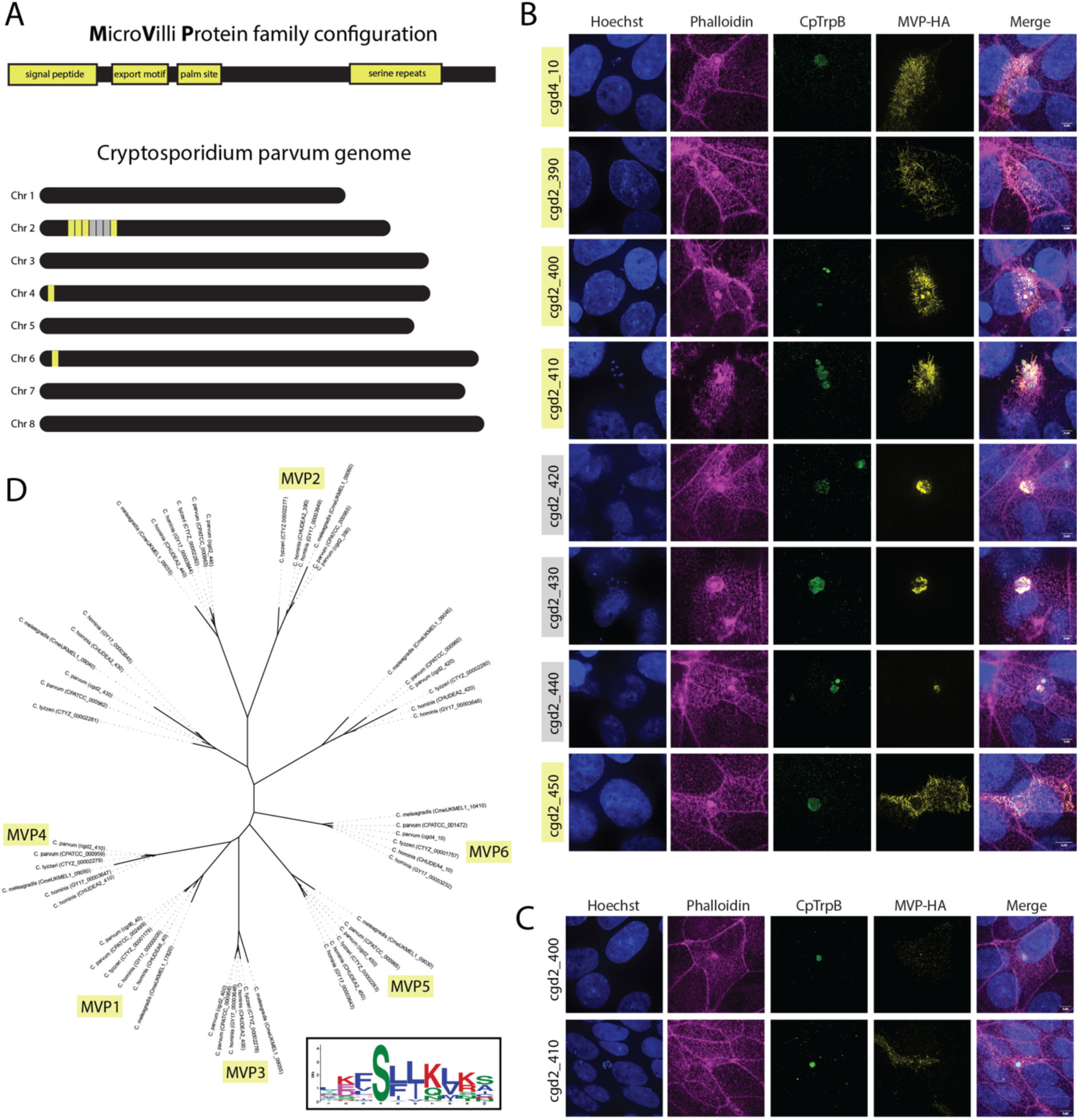
The MVP (MicroVilli Protein) family of host-exported Cryptosporidium proteins. **A)** Schematic of the shared configuration of the MVP family of *Cryptosporidium*: a signal peptide a conserved export motif, a predicted palmitoylation site, and serine repeats. Approximate location of the MVP family members on *C. parvum* chromosomes 2, 4 and 6 (yellow). **B)** IFA of epithelial cell (HCT-8) monolayer infection with wild-type *C. parvum* transfected to express epitope tagged cgd2_390 (MVP2), cgd2_400 (MVP3), cgd2_410 (MVP4), cgd2_420, cgd2_430, cgd2_440, cgd2_450 (MVP5) and cgd4_10 (MVP6). Infections were fixed at 24 hours post infection. **C)** IFA of epithelial cell monolayers infected with transgenic MVP3-HA or MVP4-HA parasites. Infections were fixed at 6 hours (MVP3-HA) and 48 hours (MVP4-HA) into infection. **D)** Phylogenetic tree of MVP family members across intestinal species of *Cryptosporidium.* Protein sequences aligned with clustal omega, then maximum-likelihood trees were constructed using FastTree with the the JTT (Jones-Taylor-Thorton) model of amino acid evolution^51^. Inset of the combined export motif from all MVP family members, constructed with the MEME suite of analysis tools^56^.

To validate these localizations and rule out an artifact of overexpression, stable lines were made with the endogenous MVP3 (cgd2_400) and MVP4 (cgd2_410) genes C-terminally tagged with the 3xHA epitope (Figure 3c). MVP3 and MVP4 were both present within sporozoites prior to host cell invasion and, similarly to MVP1, appear to export into the host cell during trophozoite stages of infection. In agreement with the transient expression experiments, both MVP3 and MVP4 localize strongly to the host cell microvilli (localisation of cgd2_410 throughout the lifecycle in Supplementary Figure 3). Further bioinformatic searches for the MVP gene family within the *Cryptosporidium* genus revealed that this family is only conserved in species of the parasite that live in the intestine of their host (Figure 3d). However, because this family is so highly polymorphic, we cannot rule out the presence of homologs in other species of *Cryptosporidium*, including gastric species. Lastly, motif based (MEME) analysis of *C. parvum* MVP family members, and homologs of related species, revealed a highly conserved putative export motif (Figure 3d inset).

### MVP1 confers virulence *in vivo* and induces the elongation of host cell microvilli

Of the MVP family, MVP1 is the most highly expressed throughout parasite replication. To answer whether MVP1 may confer virulence to *Cryptosporidium* during infection, a stable MVP1-KO line was created. Age and gender matched mice were infected with 10,000 purified and passage-matched oocysts of either the MVP1-HA or MVP1-KO transgenic C*. parvum* lines and single collections were carried out on individual mice over 21 days (Figure 4a). Parasite shedding from mice infected with MVP-KO parasites was roughly half that of mice infected with MVP1-HA. Parasites from the first passage were also purified and used to carry out a comparative infection within intestinal epithelial cell monolayers, however growth of parasites over 48 hours showed no significant difference between MVP1-HA, and MVP1-KO lines *in vitro* (Figure 4b). This suggests that MVP1 may have a function that affects parasite fitness *in vivo* but not *in vitro*.

**Figure 4.**
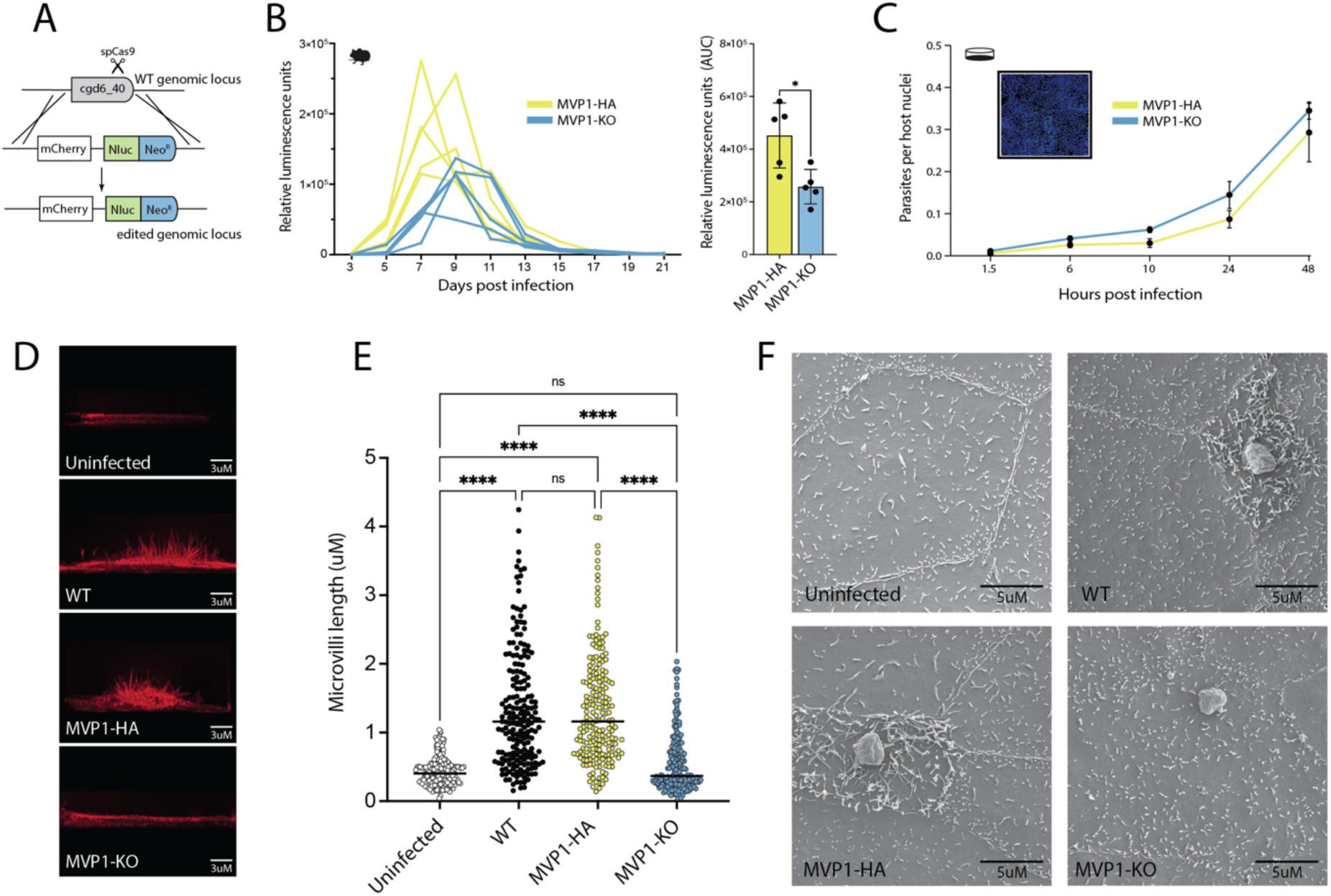
MVP1 KO affects virulence and blocks the elongation of host microvilli. **A)** Schematic of MVP1-KO genetic modification. **B)** Relative luminescence readings from collected fecal samples during infection. *Ifn*γ*^-/-^* mice were infected with either MVP1-HA (yellow) or MVP-KO (blue) parasites, where both strains contained a constitutively expressed NanoLuciferase reporter to monitor parasite burden. Mean and standard deviation for three technical repeats. Bar chart showing the area under the curve (AUC) values for each mouse infected with either MVP1-HA (yellow) or MVP1-KO (blue). Statistical analysis was done using an unpaired parametric t-test; * p ≤ 0.05. **C)** Comparative infection of epithelial cell (HCT-8) monolayers infected with either MVP1-HA or MVP1-KO parasites. Monolayers were fixed at 1.5, 6, 10, 24, and 48hrs, labelled, and then parasites were detected via microscopy (representative image inset with nuclear DNA labelled with Hoechst (blue) and parasite with Vicia villosa lectin (green)). Data points show mean and standard deviation of three biological repeats. **D)** Epithelial cells were either left uninfected or infected with MVP1-HA, wild-type, or MVP1-KO parasites for 6 hours. Cell monolayers were fixed for IFA, with host actin labelled with phalloidin. **E)** Categorical scatter plot showing measurements for the 5 longest microvilli per infected cell (40 cells / 200 measurements in total for each parasite line). Bars show mean and standard deviation. Statistical analysis was done using a one-way ANOVA; ns p>0.05, * p ≤ 0.05, ** p ≤ 0.0, *** p ≤ 0.001, **** p ≤ 0.0001. **F)** Scanning electron microscopy epithelial cell monolayers infected with MVP1-HA, MVP1-KO or wild-type *C. parvum* parasites. Monolayers were fixed at 6 hours post infection.

When imaging cell monolayers, we were surprised to discover that loss of MVP1 appeared to block the elongation of microvilli within infected epithelial cells. To test this further, monolayers of epithelial cells were infected with wild-type, MVP1-HA or MVP1-KO *C. parvum* parasites for 6 hours, then 3D rendering of infected cells from a z-series of images was used to measure host cell microvilli (Figure 4c). In epithelial cells infected with either wild-type or MVP1-HA *C. parvum,* microvilli could be seen elongated up to maximum lengths of ∼4.5μM at an average of ∼1.2μM. However, microvilli from epithelial cells infected with MVP1-KO were comparable to the uninfected controls, with an average microvilli length of ∼0.4μM (Figure 4d). This loss of elongation is even more striking when imaged by SEM, where microvilli of host cells infected with MVP1-KO parasites appear to be equivalent to that of uninfected controls (Figure 4e).

### MVP1 has evolved to function in a similar manner to the EPEC effector MAP

To determine how MVP1 might be influencing the host cell microvilli, we next sought to identify interaction partners. To accomplish this, we employed a yeast-2-hybrid screen with MVP1 as bait against a prey library created from fractionated human intestinal epithelial cell cDNA. This screen identified one highly significant interaction partner, with 70% of colonies expressing fragments of EBP50 (also known as NHERF1) (Figure 5a). EBP50 is known to be a scaffold protein, containing two PDZ domains and a C-terminal ERM (Ezrin-Radixin-Moesin) binding domain, and MVP1 appears to interact strongest with the N-terminal PDZ1 domain^28^. This interaction was supported by co-localization within epithelial cell (HCT-8) monolayers infected with MVP1-HA parasites (Figure 5b). In healthy human intestinal cells, EBP50 is known to bind a wide range of proteins at the apical membrane via its PDZ domains, facilitating the interaction of these binding partners with actin modulators. Indeed, one of the primary functions of EBP50 is the maintenance of microvilli on the apical surface of epithelial cells^29^.

**Figure 5.**
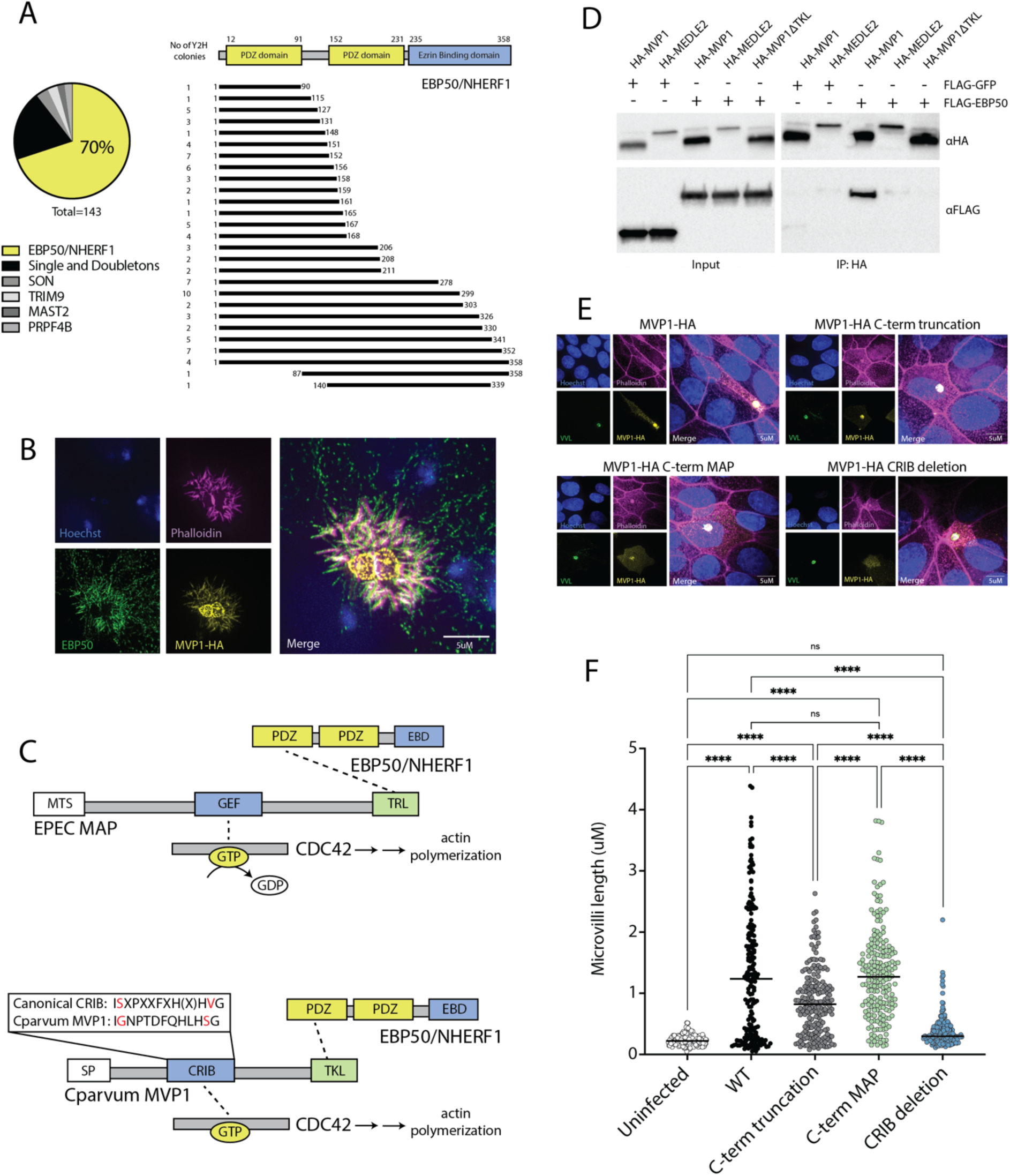
MVP1 has convergently evolved to function in a similar manner to the prokaryotic virulence factor MAP. **A)** Host interactions identified by yeast-2-hybrid using MVP1 as ‘bait’. Schematic of functional domains within EBP50 and the ‘prey’ segment identified from yeast colonies. **B)** IFA of epithelial cell (HCT-8) monolayers infected with MVP1-HA transgenic *C. parvum* parasites and co-stained for EBP50. Monolayers were fixed and labelled 6 hours post infection. **C)** Schematic of functional domains within MAP (mitochondrial targeting sequence (MTS), GEF (guanine exchange factor), and TRL (PDZ-binding domain)) and in MVP1 (signal peptide (SP), CRIB (Cdc42 and Rac interactive binding) domain, and TKL (PDZ-binding domain). **D)** HEK cells transfected to over express differing combinations of either HA-MVP1, HA-MEDLE2, or HA-MVP1ΔTKL with FLAG-GFP or FLAG-EBP50. Lysates were collected and pull downs were conducted using anti-HA beads. Eluted samples were run by western blot and probed using either an anti-HA antibody or an anti-FLAG antibody. **E)** Epithelial cells were either left uninfected or infected with wild-type, MVP1 truncation, TKL-TRL (where the last 8 amino acids of MVP1 was replaced with that of MAP), and CRIB deletion *C. parvum* parasites for 6 hours. Cell monolayers were fixed for IFA, with host actin labelled with phalloidin**. F)** Categorical scatter plot showing measurements for the 5 longest microvilli per infected cell (40 cells / 200 measurements in total for each parasite line). Bars show mean and standard deviation. Statistical analysis was done using a one-way ANOVA; ns p>0.05, * p ≤ 0.05, ** p ≤ 0.0, *** p ≤ 0.001, **** p ≤ 0.0001.

EBP50 is a known target of several bacterial effectors, including MAP – a virulence factor of the diarrheal causing bacteria enteropathogenic *Escherichia coli* (EPEC). EPEC has been shown to inject MAP into intestinal epithelial cells during infection, where this protein drives the elongation of microvilli^30,31^. MAP is known to interact with EBP50 through a C-terminal ‘TRL’ motif that binds the N-terminal PDZ domain^20,32^ (Figure 5c). *Cryptosporidium* MVP1 has a similar ‘TKL’ domain on its C-terminus and both virulence factors appear to bind the N-terminal PDZ domain of EBP50. To further test the relationship between MVP1 and EBP50, we expressed epitope tagged versions of these proteins within human embryonic kidney (HEK) cell lines (Figure 5d). From cell lysate, EBP50 coimmunoprecipitated with MVP1 but not with controls. Additionally, when the C-terminal ‘TKL’ domain was truncated, the interaction with EBP50 was lost. To take this observation back into the context of infection, we created a stable line of transgenic *Cryptosporidium* parasites lacking the last 8 amino acids of MVP1, which contains the EBP50 binding TKL domain (Figure 5e). During infection of an epithelial cell monolayer, this truncation of MVP1 resulted in a modest, but significant, decrease in host microvilli elongation (Figure 5f). This phenotype was rescued by the addition of the 8 C-terminal amino acids from EPEC MAP, which demonstrates that the EBP50 binding domains of these virulence factors function in a similar manner.

The ability of EPEC MAP to modulate host microvilli relies on the engagement of the Rho family GTPase, CDC42^31^. Further, MAP is known to interact with this key regulator of actin dynamics through an internal GEF (guanine exchange factor) domain, which binds and activates CDC42 via the exchange of GDP for GTP^32,33^ (Figure 5c). Although *Cryptosporidium* MVP1 has no recognisable GEF domains, it appears to contain a canonical CRIB, or Cdc42 and Rac interactive binding domain^34^. This interaction between MVP1 and CDC42 is also supported by *in silico* structural modelling^35^ (Supplementary Figure 4). Surprisingly, removal of this internal CRIB domain completely blocked microvilli elongation within *Cryptosporidium* infected epithelial cells (Figure 5f). This effect was independent of host-export which was not affected by the deletion of the CRIB domain (Figure 5e). Together, these observations demonstrate that the eukaryotic effector MVP1 has convergently evolved to function in an analogous manner to the prokaryotic effector MAP, engaging host EBP50 and CDC42 to drive microvilli elongation within the infected epithelial cell.

## Discussion

Apicomplexan parasites such as *Plasmodium* and *Toxoplasma* are known to export a wide repertoire of virulence factors into their infected host cells. These proteins induce diverse effects that can be cell type dependent but ultimately function to promote parasite survival and transmission^36^. Much of this export machinery appears to be conserved in *Cryptosporidium*. At the macro level, secretory organelles that house many host-exported proteins are present, including dense granules. MVP1 appears to occupy a newly discovered subset of these dense granules, called small granules, which have a separate protein content and expression pattern compared to dense granules^24^. The role of small granules during infection is still unclear, as MVP1 is the first small granule protein to be ascribed a function, but our data suggests that these organelles may be involved in host modulation, similar to dense granules. Small granules may house parasite effectors that require a different timing of production and delivery into the host cell. Within sporozoites, it was surprising to us to see a consistent localization at the apical tip of the parasite. This location is reminiscent of the apical annuli described in *Toxoplasma*, which are known sites of dense granule protein exocytosis, and it will be interesting to see if small granule proteins are transported by a similar mechanism in *Cryptosporidium*^37^.

There is also evidence of a conserved Apicomplexan export pathway at the micro level, as there are easily identifiable *Cryptosporidium* homologs to the plasmepsin family of aspartyl proteases^27^. Subsets of host-exported proteins in *Plasmodium* and *Toxoplasma* are known to contain ‘PEXEL’ and ‘TEXEL’ motifs, respectively. These motifs are recognised and cleaved by aspartyl proteases, which marks these proteins for further processing and allows for export into the infected host cell^26^. At the core of the identified MVP export motif is a serine followed by two hydrophobic residues, which bears a strong resemblance to the amino acid recognition sites for the aspartyl proteases plasmepsin IX and X^38^. The previously described *Cryptosporidium* effector MEDLE2 has a similar amino acid sequence immediately downstream of its signal peptide and, similar to MVP1, mutation of the hydrophobic residue directly following the conserved serine has shown to completely block export into the infected host cell^27^. Therefore, it is likely that MEDLE2 and the MVP family are processed and exported by a similar mechanism, but this requires more investigation. Of note, we were unsuccessful at deriving a stable line of *Cryptosporidium* with a mutated MVP1 export motif and thus could not verify cleavage of the protein, as was done for MEDLE2^27^. Because MVP1 deficient parasites are viable, we believe this failure may be due to extremely high levels of MVP1 expression, which may be lethal when the protein is unable to be properly exported.

Here, we have identified a family of six host-exported *Cryptosporidium* effectors that have conserved features and localize to the microvilli of the infected host epithelial cell. Despite these conserved features, the C-termini of the MVP family are predicted to be unstructured and share little in common with each other. Thus, we believe it is very likely that the members of the MVP family have separate and non-redundant functions within the host cell. The exception to this rule is MVP2 (cgd2_390), which has a C-terminal TKF motif, resembling the EBP50 binding motif of MVP1, TKL. Although MVP2 may be able to bind EBP50 through this domain, we observed a striking phenotype from MVP1 deficient parasites, suggesting that MVP2 may have a separate function and is unable to compensate for the loss of MVP1. Further, MVP2 also appears to lack the CRIB domain of MVP1 and may be unable to bind activated CDC42. Most of the MVP family of genes exist at a single locus at the end of the second *Cryptosporidium* chromosome. Several genes at this locus were originally predicted to be members of the MVP family (cgd2_420, cgd2_430, and cgd2_440), yet we could not detect these proteins within the host cell during infection and have chosen not include them in the MVP family. Two of these non-exported proteins, cgd2_420 and cgd2_430, have previously been described as parasite surface proteins based on antibody localization^39^. Although we cannot rule out this possibility, their location within the MVP locus is intriguing and warrants more exploration.

What is the function of microvilli elongation in *Cryptosporidium* infected cells? One possibility is that microvilli elongation is a side effect of another important function. It is crucial to state here that we did not see a difference in actin pedestal formation upon infection of epithelial cell monolayers with MVP1 deficient parasites, suggesting that microvilli elongation and actin pedestal formation may be controlled by different sets of parasite effectors and host targets. It is also important to note that, at this point, we cannot definitively state whether MVP1 drives microvilli elongation or maintains and supports microvilli elongation. Another possible function of microvilli elongation is to increase the amount of nutrients available to the parasite. Under starvation, epithelial cells in the intestine are known to elongate their microvilli to increase the apical surface area available for the uptake of sugars and other metabolites^40,41^. *Cryptosporidium* may be driving this phenotype in the absence of starvation to increase the nutrient pool available in the infected cell, which the parasite can then syphon for its own growth and replication.

Remarkably, MVP1 and MAP, effectors from eukaryotic and prokaryotic pathogens, appear to have convergently evolved to support microvilli elongation within intestinal epithelial cells through a similar mechanism. Both proteins appear to have domains that can bind CDC42 and both proteins have C-terminal domains that bind EBP50. EBP50 can be found throughout the literature under two names, EBP50 and NHERF1. EBP50 stands for Ezrin-Radixin-Moesin-binding Phosphoprotein 50, referring to this protein’s ability to bind actin cytoskeleton modulators, such as ezrin, that are required for maintenance of intestinal microvilli^42^. NHERF1 stands for Na/H Exchanger Regulatory Factor 1, which refers to another role of this protein, influencing sodium/hydroxide exchange (predominantly through NHE3) and therefore water homeostasis in the intestine^43^. The *Cryptosporidium* parasite causes a diarrheal disease, yet the underlying process that causes these symptoms is poorly understood. Other intestinal pathogens, such as EHEC and Vibrio cholerae, are known to drive diarrheal symptoms through disruption of NHE3^44,45^. MVP1’s interaction with EBP50/NHERF1 may be one of the key underlying mechanisms that drive the diarrheal symptoms of cryptosporidiosis, and this interaction deserves more attention.

## Methods

### Mouse models of infection

Ifnγ^-/-^ mice were bred in the Francis Crick Institute Biological Research Facility under specific pathogen free conditions. Age and gender matched mice in this study range from 4 to 6 weeks for comparative experiments. Both male and female Ifnγ^-/-^ mice, age 4-12 weeks were used for generation and propagation of *C. parvum* transgenic parasite lines, with no notable difference in parasite burden or shedding. Experiments were undertaken in accordance with UK Home Office regulations under project license PP8575470.

### Cell lines

Human ileocecal adenocarcinoma (HCT-8) cells were purchased from ATCC, these were mycoplasma tested at The Francis Crick Institute and when determined negative for mycoplasma were used for experiments.

### Parasite strains

Wild-type *C. parvum* was purchased from Bunchgrass Farms (Dreary, ID) to be used in experiments and to create all *C. parvum* transgenic lines which were made and propagated in *Ifn*γ*^-/-^* mice. Transgenic lines were isolated from faecal samples collected daily using sucrose floatation and subsequent caesium chloride density gradient as previously described^46^.

### Generation of transgenic parasites

Transgenic *C. parvum* parasites were created using previously described methods^47,48^. 2.5×10^7^ parasites were bleach-treated in 1% sodium hypochlorite (VWR, 301696S), washed in 1%FBS/Roswell Park Memorial Institute (RPMI) (Gibco, A1049101) media and resuspended in 0.75% sodium taurocholate (Merck, 86339) for 1 hour to artificially induce excystation of oocysts. Excysted sporozoites were transfected with a repair template and a plasmid expressing Cas9 and a guide targeting various locations within the *C. parvum* genome, Nanoluciferase and a Neomycin resistance cassette. Transfected parasites at this stage were used either to infect HCT-8 cell monolayers in culture or mice. Mice were orally gavaged first with saturated sodium bicarbonate (Sigma, S5761) and then the transfected sporozoites. Following infection mice were treated with paromomycin (16g/L) (Biosynth Ltd, AP31110) within drinking water allowing for selection of transfected parasites and parasite burden was measured using a nanoluciferase assay (as described below).

### Nanoluciferase assay to monitor parasite shedding

20mg of faecal material was added to 3mm glass beads and dissolved using lysis buffer. Following vortexing at 2000 rpm for 15 minutes at room temperature (RT) this was combined 1:1 with nanoluciferase substrate/nanoluciferase assay buffer (1:50) (Promega, N1150) in a white bottom plate and relative luminescence was determined using a BioTek citation 5.

### *In vitro*infection and immunofluorescence assay

Coverslips were seeded with HCT-8 cells in 10%FBS/RPMI media and left to grow to 70-80% confluency. Wild-type or transgenic *C. parvum* oocysts were bleached, washed in 1%FBS/RPMI media, and resuspended in 0.75% sodium taurocholate for 10 minutes. Sporozoites were centrifuged at 16,000xg for 3 minutes, resuspended in 1%FBS/RPMI and added to HCT-8 monolayers. Cells were fixed with 4%PFA/PBS at various timepoints into infection, permeabilized with 0.1%Triton-X-100/PBS and blocked in 4%BSA/PBS. Antibodies were diluted in 1%FBS/PBS. The primary antibodies rat monoclonal anti-HA (Roche, 11867423001) and rabbit anti-trpB antibody were used at 1:1000. The Alexa Fluor 594 donkey anti-rat (Life technologies, A-21209) and Alexa Fluor 488 goat anti-rabbit (Thermofisher, A32731) secondary antibodies were used at 1:1000. Alexa Fluor 647 Phalloidin (Thermofisher, A22287) was added alongside secondary antibody at 1:1000 and Hoechst 33342 (Invitrogen, H3570) at 1:10,000. Coverslips were mounted onto slides using Prolong Gold antifade (Thermofisher, P36934) and imaged using a VT-iSIM microscope.

### Permeabilzation assay

Coverslips confluent with HCT-8 cells were infected with wild-type or MVP-HA transgenic *C. parvum* and stained as described above. Non-permeabilized cells were fixed with 1%PFA/PBS and were not permeabilized with 0.1%Triton-X-100/PBS before blocking.

### Immunohistochemistry on infected intestine

Infected *Ifn*γ*^-/-^* mice were euthanized between day 10-12 of infection and 7cm of ileum was dissected. Tissue was flushed with saline and fixed for 48 hours in 10% formalin. For protection during cryosectioning the tissue section was then placed in 30%sucrose/PBS, mounted into gelatine and frozen. Blocks were sectioned at 5-micron thickness. Tissue sections were permeabilized, blocked and stained as described above.

### Expansion microscopy of sporozoites and IFA

(Protocol adapted from Liffner and Absalon, 2021^49^) Coverslips within a 24-well plate were coated with poly-D-lysine for 1 hour at RT before being washed twice with PBS. Newly excysted sporozoites in 1%FBS/RPMI were added to adhere for 15 minutes at 37°C. RPMI was removed and coverslips incubated for 15 minutes with 4%PFA/PBS before being washed twice with PBS. Protein crosslinking prevention was performed by mixing a formaldehyde/acrylamide/PBS solution and adding this to wells. Plates were incubated overnight at 37°C. Gelation was performed by pre-cooling a humid chamber at -80°C for 10 minutes before placing on ice. Coverslips were washed with PBS, blotted dry and placed top down onto gel mounting slides. Working quickly TEMED and then APS (after being thawed on ice for 30 minutes) were added to monomer solution (sodium acrylamide/acrylamide/BIS/PBS) vortexed and pipetted under coverslips. Coverslips were incubated on ice before being moved to a 37°C incubator for 1 hour.

Coverslips were moved into a 6-well plate with denaturation buffer and incubated at room temperature with agitation for 15 minutes. Gels were then separated from the coverslips and moved into Eppendorf’s filled with fresh denaturation buffer followed by incubation at 95oC for 1h30minutes. Gels were moved into petri dishes with dH20 and incubated at room temperature with agitation for 30 minutes. Petri dishes were replaced with fresh dH20 and this was repeated for a total of 3 dH20 changes. dH20 was replaced with PBS for 15minutes and then once more with fresh PBS. Gels were transferred into a 6 well plate with 2%BSA/PBS and incubated with agitation at room temperature for 1 hour. The 2%BSA/PBS was replaced with primary antibody in 2%BSA/PBS and incubated at room temperature overnight. Gels were washed 3x with 0.5%Tween20/PBS for 10 minutes with agitation before being incubated with secondary antibody/Hoescht/NHS ester in PBS for 2-3 hours at room temperature with agitation. Gels were washed 3x with 0.5%Tween20/PBS for 10 minutes with agitation. Gels were moved into petri dishes with dH20 and incubated at room temperature with agitation for 30 minutes. dH20 was replaced with fresh dH20 for a total of 3 dH20 changes. Gel size was measured to calculate expansion factor. Poly-d-lysine was added to a 60mm dish and incubated for 1 hour to overnight before being washed 3x with dH20. Gels were transferred into dishes and imaged using a VT-iSIM microscope.

### BFA treatment during *C. parvum* infection

Coverslips were seeded with HCT-8 cells in 10%FBS/RPMI media left to grow to 70-80% confluency. Wild-type or transgenic *C. parvum* oocysts were bleached and washed in 1%FBS/RPMI media before being resuspended in 0.75% sodium taurocholate for 10 minutes. Parasites were then centrifuged at 16,000xg for 3 minutes and resuspended in 1%FBS/RPMI to be used to infect HCT-8 monolayers in 1%FBS/RPMI media. Cells were either fixed at 10 hours into infection with 4%PFA/PBS or were treated with 10μg/ml Brefeldin-A (Sigma, B5936)/1% FBS/RPMI at 3 hours into infection before being fixed with 4% PFA/PBS at 10 hours. Coverslips were then permeabilized with 0.1%Triton-x-100, blocked with 4% BSA/PBS and stained for IFA.

### Western blot on *C. parvum* infected cells

For sporozoite lysate, 1×10^7^ wild-type *C. parvum* oocysts were bleached, washed in 1%FBS/RPMI media, and resuspended in 0.75% sodium taurocholate for 1 hour. Sporozoites were pelleted and lysed in 1% RIPA buffer supplemented with both protease inhibitor cocktail and 150μM PMSF for 10 minutes. For cell cultures, HCT-8 cells were infected with 8×10^5^ oocysts for 24 or 48 hours, washed with ice cold PBS and lysed in 1% RIPA buffer supplemented with both protease inhibitor cocktail and 150μM PMSF for 10 minutes. HCT-8 cells were scraped, and lysate was collected. Samples were sonicated with a microtip sonicator on ice for 3 rounds of 30 seconds at an amplitude of 30% and insoluble and soluble fractions were separated by centrifugation at 4°C at 16,000xg for 20 minutes. Samples were mixed with 5x non-reducing sample buffer (pierce) and boiled at 95°C for 10 minutes, loaded into a 12% Mini-PROTEAN TGX Precast Protein Gel (Bio-Rad) and ran at 90v for 1 hour. Gels were transferred onto a 0.2 μM Nitrocellulose membrane (Bio-Rad) and the membrane was blocked using 3%Milk/ 0.01%Tween-20/PBS. Rat monoclonal anti-HA (Roche) was diluted in blocking solution 1:1000 and added for 1 hour. Membranes were washed 3x with 0.01%Tween-20/PBS and HA Tag polyclonal HRP antibody was added 1:1000 diluted in blocking solution for 1 hour. Membranes were washed with 0.01%Tween-20/PBS 3x for 10 minutes. Clarity Western ECL substrate was added to membranes which were then imaged using a ChemiDoc Imaging System (BioRad).

### Yeast-two-hybrid screen

The Y2H screening was carried out by Hybrigenics Services SAS as previously described in Guérin *et al.,* 2017^50^ using a colon tumour epithelial cell cDNA library (a combination of HCA7, Caco2, Colo205 and SW480 cells). The bait construct used was full length MVP with the signal peptide removed.

### Microvilli length Quantifications

Transgenic *C. parvum* oocysts were excysted and used to infect coverslips confluent with HCT-8 cells as above. Coverslips were fixed with 4%PFA/PBS at 6 hours into infection, permeabilized with 0.1%Triton-x-100/PBS and blocked in 4%BSA/PBS. Antibodies and stains were diluted in 1%FBS/PBS. Alexa Fluor 647 Phalloidin was added at 1:1000, VVLectin fluorescein (2B Scientific, FL-1231-2) was added at 1:4000 and Hoechst 33342 (Invitrogen, H3570) at 1:10,000. Coverslips were blinded externally and mounted onto slides using Prolong Gold antifade (Thermofisher, P36934). Z stack images were taken using a VT-iSIM microscope and rendered as a 3D image at 90° using Huygens essential. Using ImageJ, the 5 longest microvilli viewed by Alexa Fluor 647 Phalloidin (Thermofisher, A22287) on the apical surface of 40 HCT-8 cells per *C. parvum* strain were measured as a proxy for microvilli length.

### Preparation of cell monolayers for SEM

HCT-8 monolayers were cultures on coverslips and infected as above for 6 hours. Coverslips were washed with warm 1%RPMI and left in 500ul fresh 1%RPMI. Coverslips were then fixed at room temperature by adding to the medium a pre-warmed (37°C) double-strength fixative - 8% Formaldehyde in 0.2 M phosphate buffer - for 15 minutes at a 1:1 ratio. Fixative was removed and replaced with 2.5%Glutaldehyde/4%Formaldehyde in 0.1M phosphate buffer for 30 minutes at room temperature. Coverslips were washed 3x for 5 minutes with 2ml of 0.1M phosphate buffer. Next coverslips were covered with 1% OsO4/H20 and kept at 4°C for 1 hour. Coverslips were washed 3x with Deuterium-depleted water before being dehydrated with a graded ethanol series: 2ml of 50%, 70%, 90% and 100% (2x) ethanol for 5 minutes each step at room temperature. Samples were dried using a Critical Point Dryer 300 in ethanol (programme 03 – human cells). Stubs were mounted with double-faced carbon tape using a Quorum Q150 using the programme Platinum 5nm slow. Imaging of samples was carried out on an SEM Quanta 250 (3kV, spot 2, WD 9.5mm, High vacuum, dwell time: 10 µs, store resolution 3k).

### Phylogenetic tree analysis

Protein sequences were aligned with clustal omega (version 1.2.0), maximum-likelihood trees were constructed using FastTree (version 2.1.11) with the JTT (Jones-Taylor-Thorton) model of amino acid evolution and trees were visualized using iTOL (version 7)^51–54^.

### MEME plots

The MEME suite tool Multiple Em for Motif Elicitation (MEME version 5.5.7) was used in classic mode with default parameters to scan the N-terminus of the MVP and MEDLE protein families to discover motifs (MVP1-6, cgd6_5490, cgd6_5480, cgd5_4600, cgd5_4580, cgd5_4610 and cgd5_4590 from C. parvum, C. hominis and C. tyzzeri isolates)^55^.

### Co-immunoprecipitations

HEK293ET (CVCL_6996) cells were seeded in 10% FBS/RPMI supplemented with Amphotericin B (250 μg/ml), Penicillin (100 units/ml) and Streptomycin (100 μg/ml). Cells were transfected with pcDNA4/TO-HA-MVP1, pCDNA4/TO-HA-GFP, pcDNA4/TO-MVP11TKL or pCDNA4/TO-HA-MEDLE2 along with pCDNA4/TO-EBP50 or pCDNA4/TO-GFP 18 hours after seeding using Lipofectamine 3000 (ThermoFisher) following the manufacturers protocol. Twenty-four hours post-transfection HEK293ET cells were washed in PBS and lysed in RIPA buffer (50 mM Tris, pH 8.0, 150 mM NaCl, 0.5 M EDTA, 1% NP-40, 0.5% sodium deoxycholate, 0.1% SDS supplemented with cOmplete™, Mini, EDTA-free Protease Inhibitor Cocktail (Roche)) on ice. Lysates were incubated on ice for 15 min and vortexed intermittently before centrifugation at 17000xg for 15 minutes at 4°C. An aliquot was taken to probe input, before lysates were diluted 10-fold in dilution buffer (50 mM Tris, pH 8.0, 150 mM NaCl, 1% NP-40). Samples were incubated on a rotor overnight with monoclonal anti-HA−Agarose (Merck, A2095). After 18 hours of incubation anti-HA-agarose was washed in dilution buffer and pelleted by centrifugation at 5000xg, this was repeated 3 times. After the final wash the anti-HA-agarose was centrifuged at 17000xg, before being incubated in 4× sample loading dye (0.5 M Tris, pH 6.8, 40% glycerol, 6% SDS, 1% bromophenol blue, and 0.8% β-mercaptoethanol) and analysed by immunoblotting. Antibodies used were the anti-FLAG antibody (Merck, F7425) and the anti-HA-tag antibody (Cell Signalling Technology, 3724S).

## Acknowledgements

We would like to thank the Scientific Technology Platforms (STPs) at The Francis Crick Institute, particularly the Advanced Light Microscopy for providing equipment and training and the Biological Research Facility for breeding and maintenance of mouse lines. We would like to thank VEuPathDB for maintenance of critical databases that were essential for this work. We would also like to thank Dr Francesca Ester Morreale for her valuable insight. This work was supported by The Francis Crick Institute, which receives its core funding (CC2063) from Cancer Research UK, the Medical Research Council, and the Wellcome Trust, as well as a grant from UKRI (101042783).

## Author Contributions

ER and AS developed the concept and outline of the study. Experiments were designed and executed by ER, MP, LCS, KS, and AS. Bioinformatics expertise and training was provided by TTM. ER and AS drafted the manuscript with input from all listed authors.

---

**Supplementary Figure S1.**
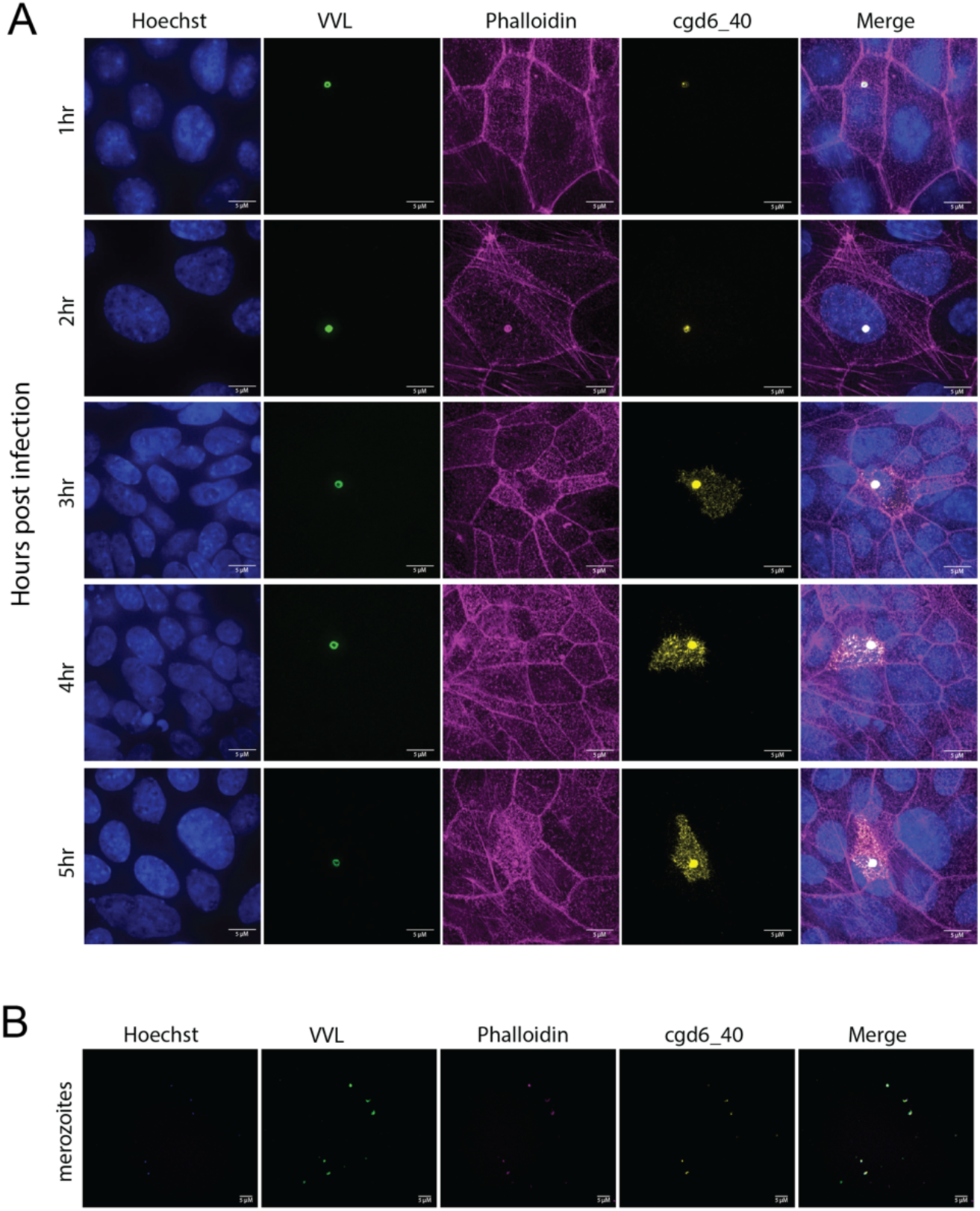
Host-export of cgd6_40 following invasion. **A)** IFA of epithelial cell (HCT-8) monolayers infected with transgenic cgd6_40-HA parasites. Infections were fixed at one-hour intervals following infection. **B)** Isolated cgd6-40-HA merozoites demonstrating cgd6_40 expression prior to reinvasion.

**Supplementary Figure S2.**
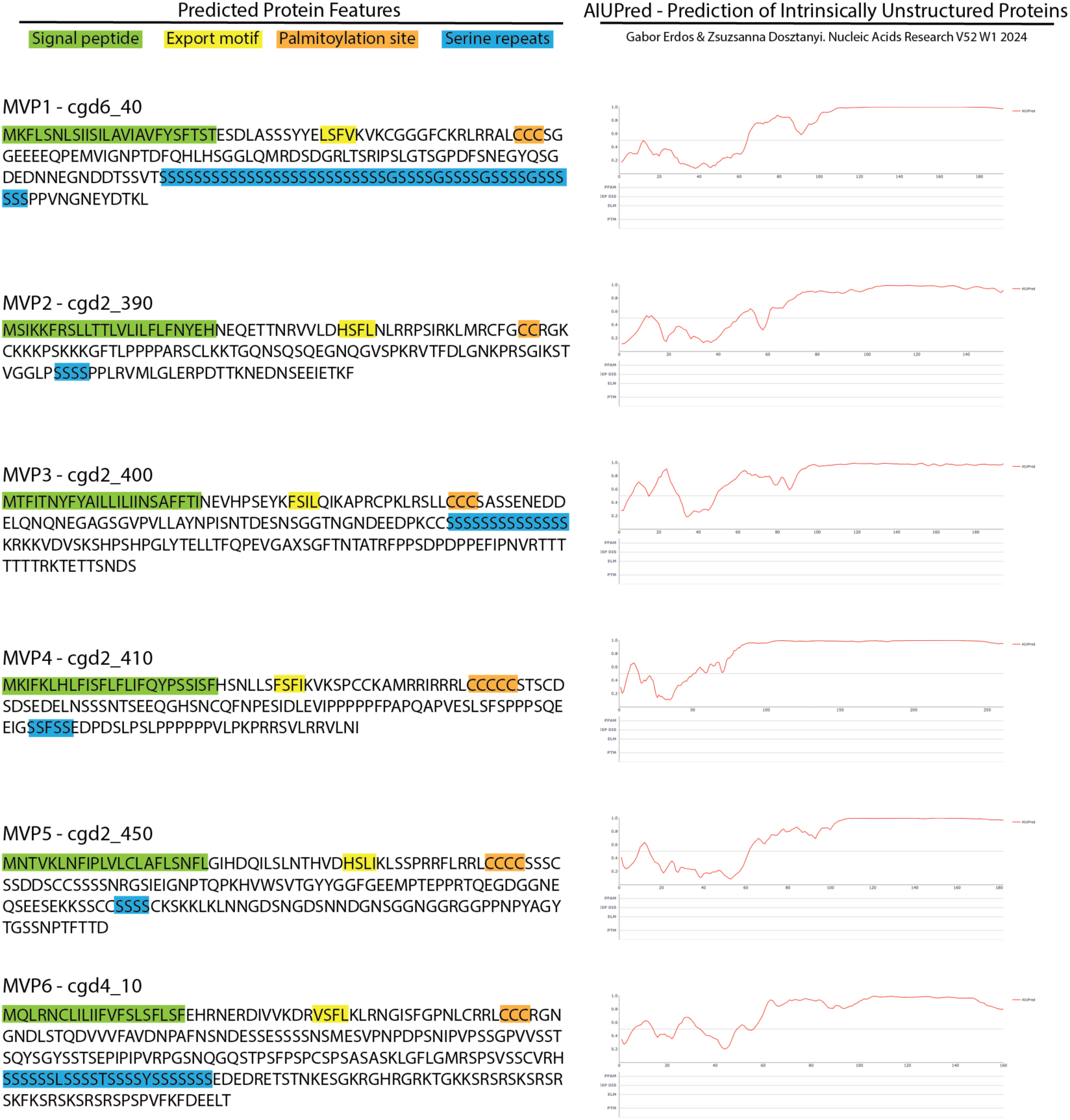
Protein feature of the Cryptosporidium parvum MicroVilli Protein family. Predicted features on left with signal peptide prediction via CryptoDB^22^. Prediction of intrinsic disorder on right via AIUPred^57^.

**Supplementary Figure S3.**
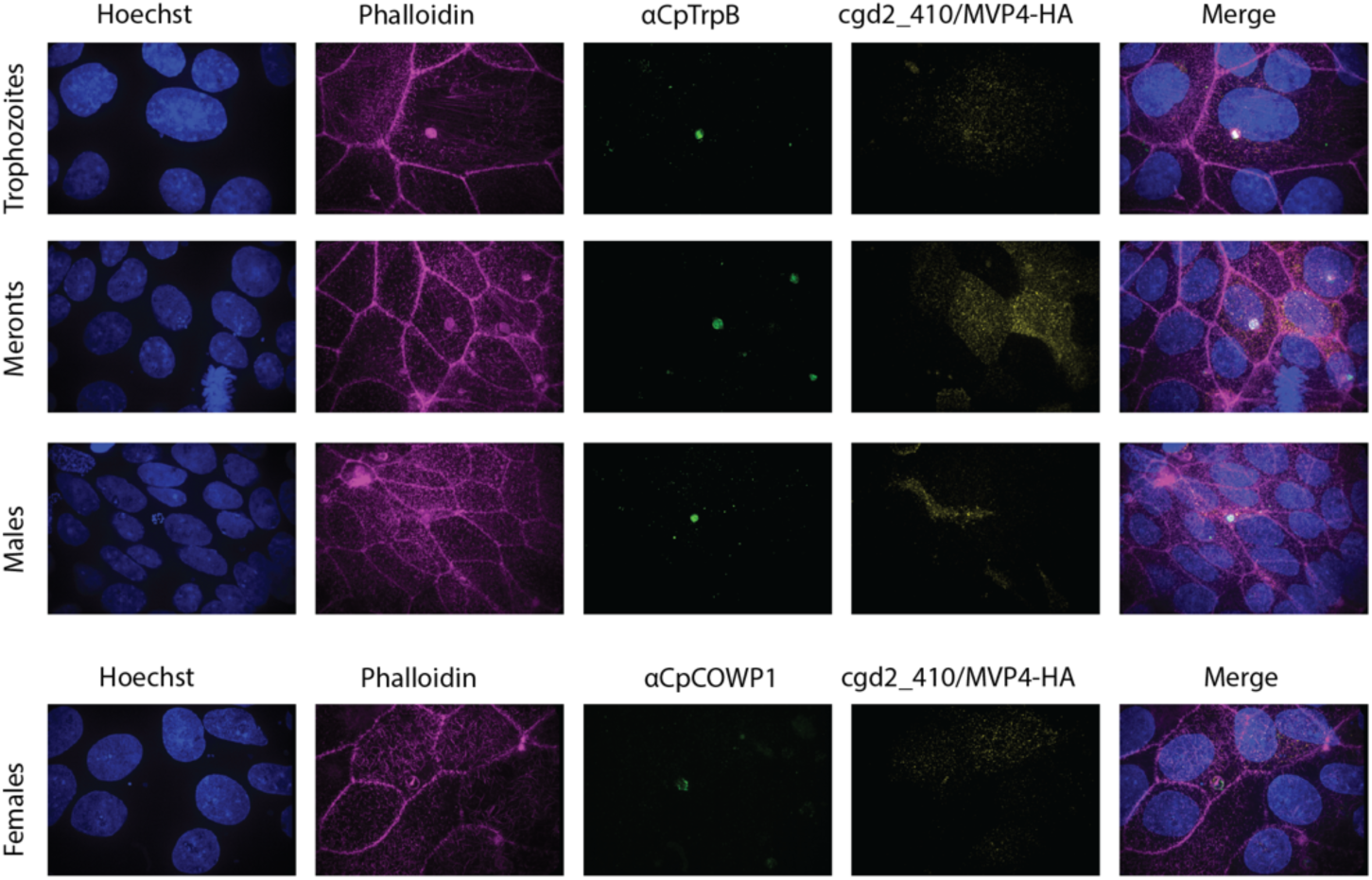
Lifecycle expression and localisation of cgd2_410-HA. IFA of epithelial cell (HCT-8) monolayers infected with transgenic cgd6_40-HA parasites. Infections were fixed at 6 hours (trophozoites), 24 hours (meronts) and 48 hours (males and females).

**Supplementary Figure S4.**
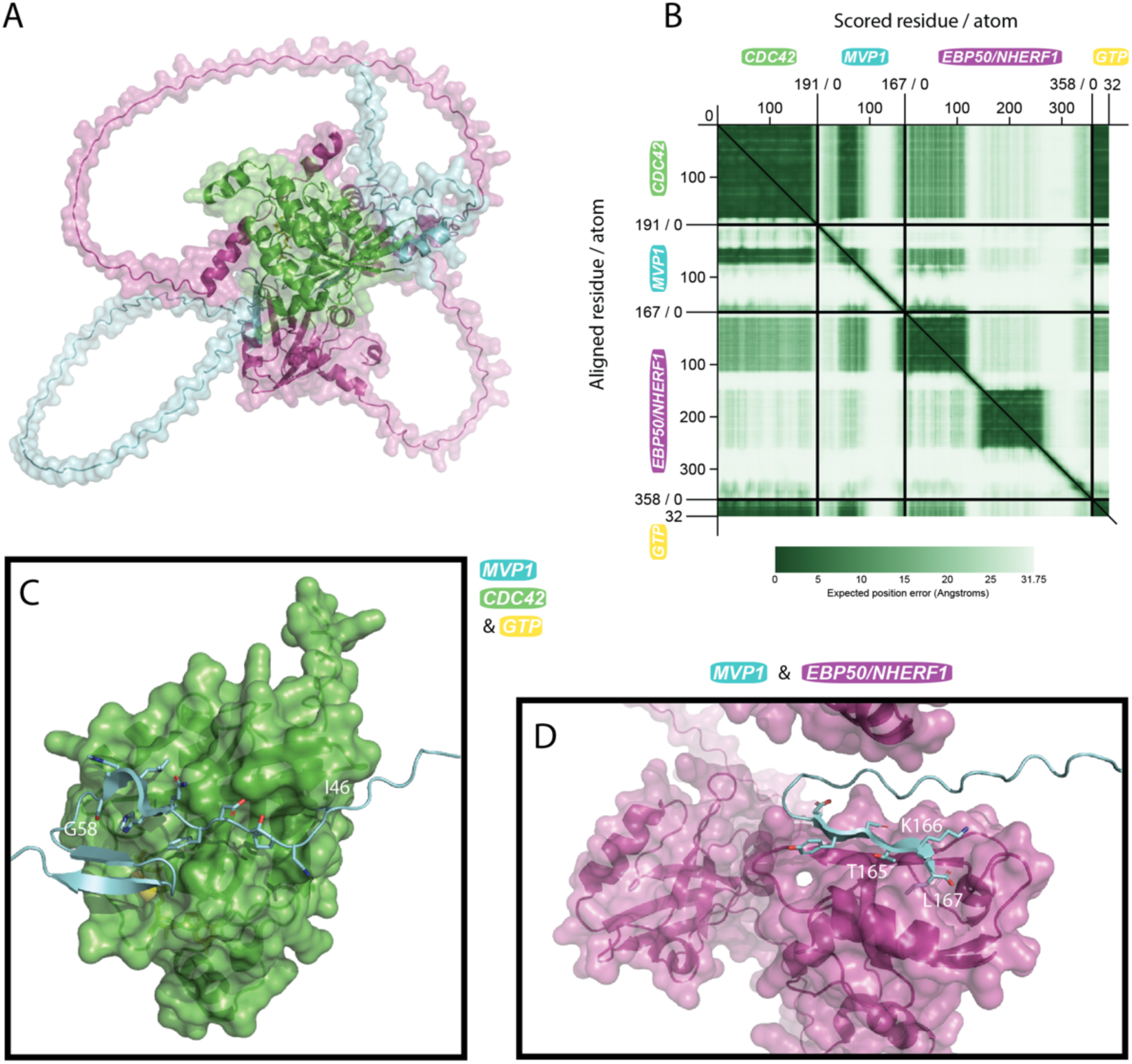
Prediction of MVP1 interactions with EBP50 and CDC42. Amino acid sequences of MVP1 (cgd6_40 minus signal peptide), human EBP50 (UniProt O14745), human CDC42 (UniProt P60953), and ligand GTP were used for modelling via AlphaFold^35^. Structure visualised via PyMol and PAEviewer^58^.

